# Mapping localization of 21 endogenous proteins in the Golgi apparatus of rodent neurons

**DOI:** 10.1101/2022.11.30.518582

**Authors:** Danique M van Bommel, Ruud F Toonen, Matthijs Verhage

## Abstract

The Golgi apparatus is the major sorting hub in the secretory pathway and particularly important for protein sorting in neurons. Knowledge about protein localization in Golgi compartments is largely based on work in cell lines. Here, we systematically compared protein localization of 21 endogenous proteins in the Golgi apparatus of mouse neurons using confocal microscopy and line scan analysis. We localized these proteins by measuring the distance relative to the canonical TGN marker TGN38. Based on this, proteins fell into three groups: upstream of, overlapping with or downstream of TGN38. Seven proteins showed complete overlap with TGN38, while proteins downstream of TGN38 were located at varying distances from TGN38. Proteins upstream of TGN38 were localized in between TGN38 and the *cis*-/medial Golgi markers Giantin and GM130. This localization was consistent with protein function. Our data provide an overview of the relative localization of endogenous proteins in the Golgi of primary mouse neurons.

## Introduction

The Golgi apparatus consists of *cis*-, medial and *trans*-Golgi cisternae that continue into a tubular *trans*-Golgi network (TGN). Resident Golgi proteins are distributed along the Golgi stack to enable diverse functions including post-translational modifications, such as glycosylation, of cargo proteins. Newly synthesized cargo proteins arrive at the *cis*-Golgi and are processed as they progress to the TGN, where they are sorted for export to diverse organelles or for secretion. In neurons, secretion occurs via constitutive and regulated secretory pathways. The regulated secretory pathway contains neuropeptides and neurotrophins that modulate synaptic activity and thereby regulate diverse functions such as memory, fear and appetite [1–3]. Experiments to study protein localization in the Golgi apparatus are often performed in cell lines. The observed localization might not be the same as in cells with a regulated secretory pathway, such as neurons. In addition, previous studies have combined data from antibody staining and overexpression experiments [4–6], but overexpression of proteins might affect their localization and can even induce perturbations of the Golgi [7,8]. Finally, most studies focus on the localization of one or a few proteins.

Here, we studied the relative localization of 21 endogenous proteins in the Golgi apparatus of primary mouse neurons using high-resolution confocal microscopy of well separated Golgi stacks. While protein localization was consistent with their proposed functions, we identify a relatively large number of proteins localized further downstream than the canonical TGN marker TGN38.

## Results

We used confocal microscopy to characterize the relative localization of endogenous proteins in the Golgi apparatus of mouse neurons. Combining a *cis*-(/medial) Golgi marker with a TGN marker allowed us to identify areas of the Golgi apparatus with well-separated Golgi-stacks (Fig. 1a,b). By drawing a line through these stacks and plotting intensity profiles along the line, we determined the localization of proteins relative to the *cis*-(/medial) Golgi and TGN (Fig. 1c). Since Golgi stacks are randomly oriented within the three dimensions of a cell, some bias was introduced by selecting sections of the Golgi apparatus that were in the plane of view. As a result, it was impossible to measure absolute distances between proteins and we chose to describe the localization of proteins relative to each other. To prevent further bias, the lines were drawn based on *cis-*/medial Golgi and TGN staining, without visualizing the protein of interest. Applying this approach to confocal images resulted in similar information as STED images (Sup. Fig. 1). We used this tool to analyze the relative localization of 21 proteins. We selected several proteins that are commonly used as Golgi markers (e.g. giantin and GM130), proteins that play a role in regulated secretion (e.g. HID1 and CCDC186) and proteins of which endogenous localization has not been studied before (e.g. CASC4 and PRR36). These proteins show high staining intensity in the Golgi, but are not necessarily exclusively localized to the Golgi apparatus. The analyzed proteins could be categorized into three distribution groups based on their localization along the Golgi axis relative to TGN38: proteins with an intensity peak upstream of TGN38 (i.e. *cis*-/medial Golgi), overlapping with TGN38 (i.e. TGN) and downstream of TGN38. We chose to plot protein distribution relative to TGN38 because the antibody against this protein could be combined with any antibody species for the protein of interest.

**Fig. 1.**
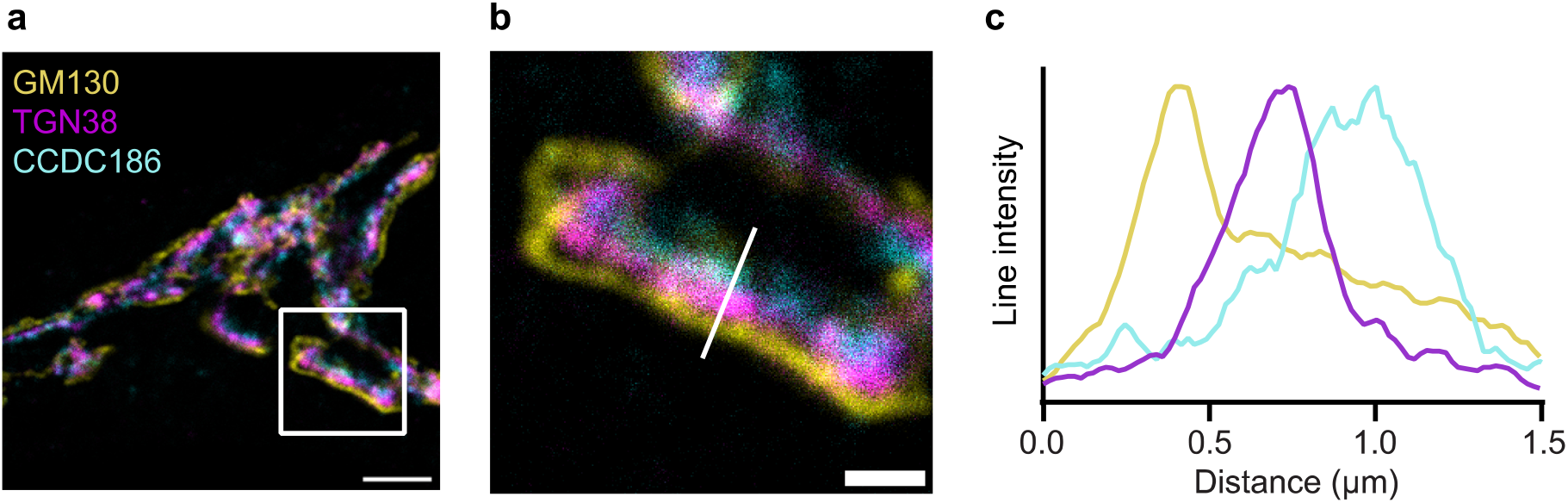
Golgi stacks visualized using confocal microscopy. **a** Neuron stained for GM130 (*cis*-Golgi), TGN38 (TGN) and CCDC186. **b** Zoom from the white square area marked in **a. c** Line intensity profile along white line in **b**. Intensities were normalized and smoothed before plotting. Scale bar is 3 μm (**a**) and 1 μm (**b**).

### Proteins downstream of TGN38

For every neuron, *cis*-/medial Golgi and TGN were clearly disthinghuisable (Fig. 2a (See also Fig. 3a, Fig. 4a, Sup. Fig. 3a, Sup. Fig. 4a)). While AP1, AP3, ARFIP2 and CCDC186 localized near TGN38 and GM130/giantin, no overlap was detected (Fig. 2b,c). Zooms showed that for all four proteins, three layers were detected: *cis*-/medial Golgi, TGN and proteins of interest (Fig. 2d). Distance from peaks of intensity of TGN38 was larger than 0 for AP1, AP3, ARFIP2 and CCDC186 and distance from giantin or GM130 was even larger (Fig. 2e). In some cells, ARFIP2 overlapped more with TGN38 (Sup. Fig. 2). Supplementary figure 3 shows that Stx16, Stx6, Vti1a (C-terminus), VAMP4 and tomosyn were also localized downstream of TGN38. The distance between TGN38 and those proteins was smaller than the distance between TGN38 and AP1, AP3, ARFIP2 or CCDC186. Altogether, AP1, AP3, ARFIP2, CCDC186, Stx16, Stx6, Vti1a (C-terminus), VAMP4 and tomosyn localized downstream of TGN38.

**Fig. 2.**
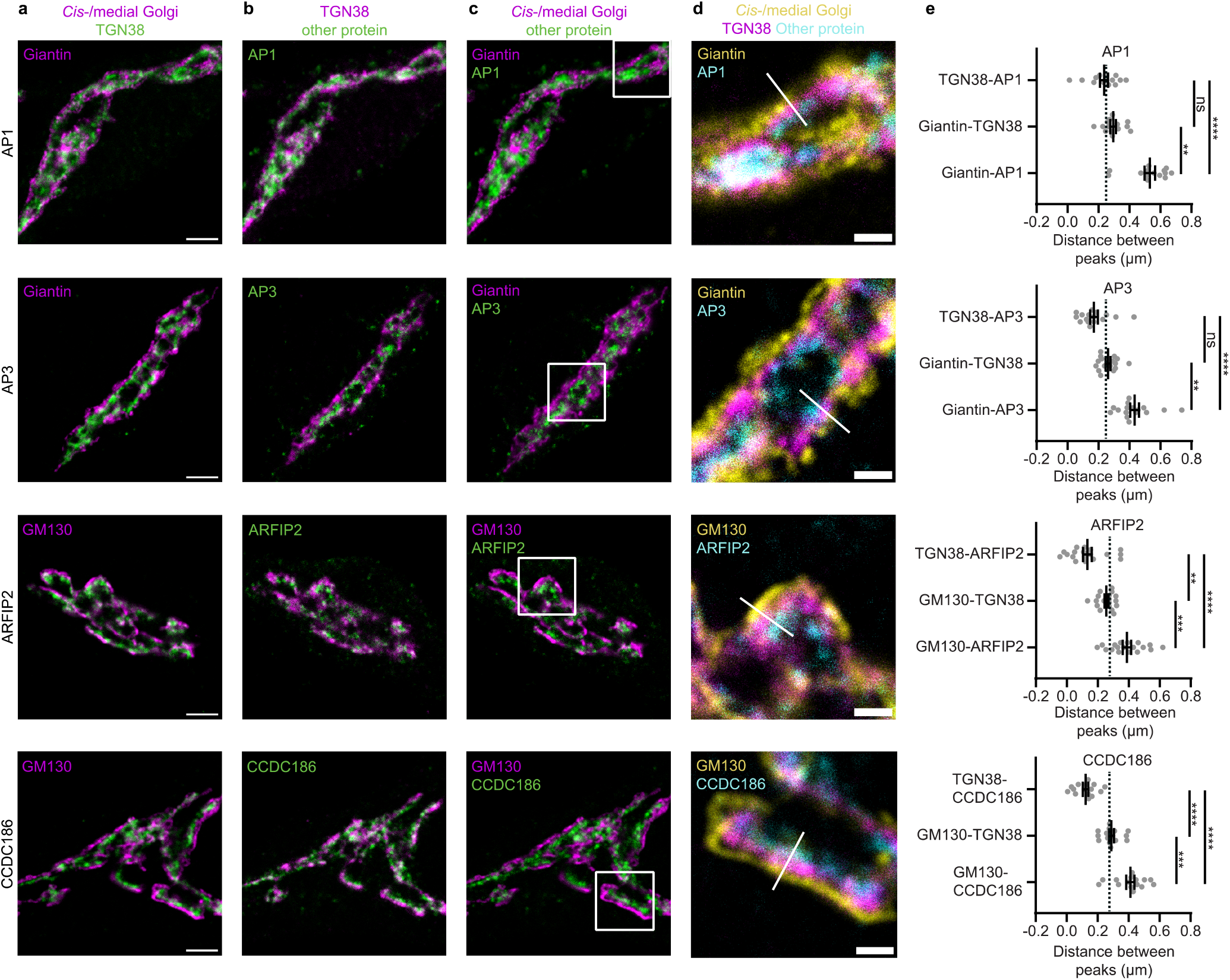
Several Golgi proteins localize downstream of TGN38. **a-c** Representative examples of neurons immunostained for a *cis*-/medial Golgi marker (GM130 or giantin), TGN38 and a protein of interest (AP1, AP3, ARFIP2 and CCDC186). Images showing immunostaining of *cis*-/medial Golgi marker and TGN (**a**), TGN and protein of interest (**b**), *cis*-/medial Golgi and protein of interest (**c**). **d** Zoom from the white square area marked in **c**, showing all three stained proteins. White line is a representative example of a line drawn for intensity plotting. **e** Distance between intensity peaks of TGN38 and protein of interest, *cis*-/medial Golgi and TGN; and *cis*-/medial Golgi and protein of interest, measured by line scans. AP1: (TGN38-AP1: n = 14; Giantin-TGN38: n = 14; Giantin-AP1: n = 14). Kruskall-Wallis test with Dunn’s multiple comparisons test: TGN38-AP1 vs Giantin-TGN38: non-significant (ns), TGN38-AP1 vs Giantin-AP1: ****p < 0.0001, Giantin-TGN38 vs Giantin-AP1: **p = 0.0036. AP3: (TGN38-AP3: n = 16; Giantin-TGN38: n = 16; Giantin-AP3: n = 16). Kruskall-Wallis test with Dunn’s multiple comparisons test: TGN38-AP3 vs Giantin-TGN38: non-significant (ns), TGN38-AP3 vs Giantin-AP3: ****p < 0.0001, Giantin-TGN38 vs Giantin-AP3: **p = 0.0042. ARFIP2: (TGN38-ARFIP2: n = 18; GM130-TGN38: n = 18; GM130-ARFIP2: n = 18). One-way ANOVA with Tukey’s multiple comparisons test: TGN38-ARFIP2 vs GM130-TGN38: **p = 0.0018, TGN38-ARFIP2 vs GM130-ARFIP2: ****p < 0.0001, GM130-TGN38 vs GM130-ARFIP2: ***p = 0.0009. CCDC186: (TGN38-CCDC186: n = 15; GM130-TGN38: n = 15; GM130-CCDC186: n = 15). One-way ANOVA with Tukey’s multiple comparisons test: TGN38-CCDC186 vs GM130-TGN38 and TGN38-CCDC186 vs GM130-CCDC186: ****p < 0.0001, GM130-TGN38 vs GM130-CCDC186: ***p = 0.0004. Dotted lines represent average distance between TGN38 and GM130 or TGN38 and giantin in the entire dataset. Bars show mean ± SEM. Detailed statistics are shown in Supplementary Table 1. Scale bar is 3 μm (**a**) and 1 μm (**d**).

**Fig. 3.**
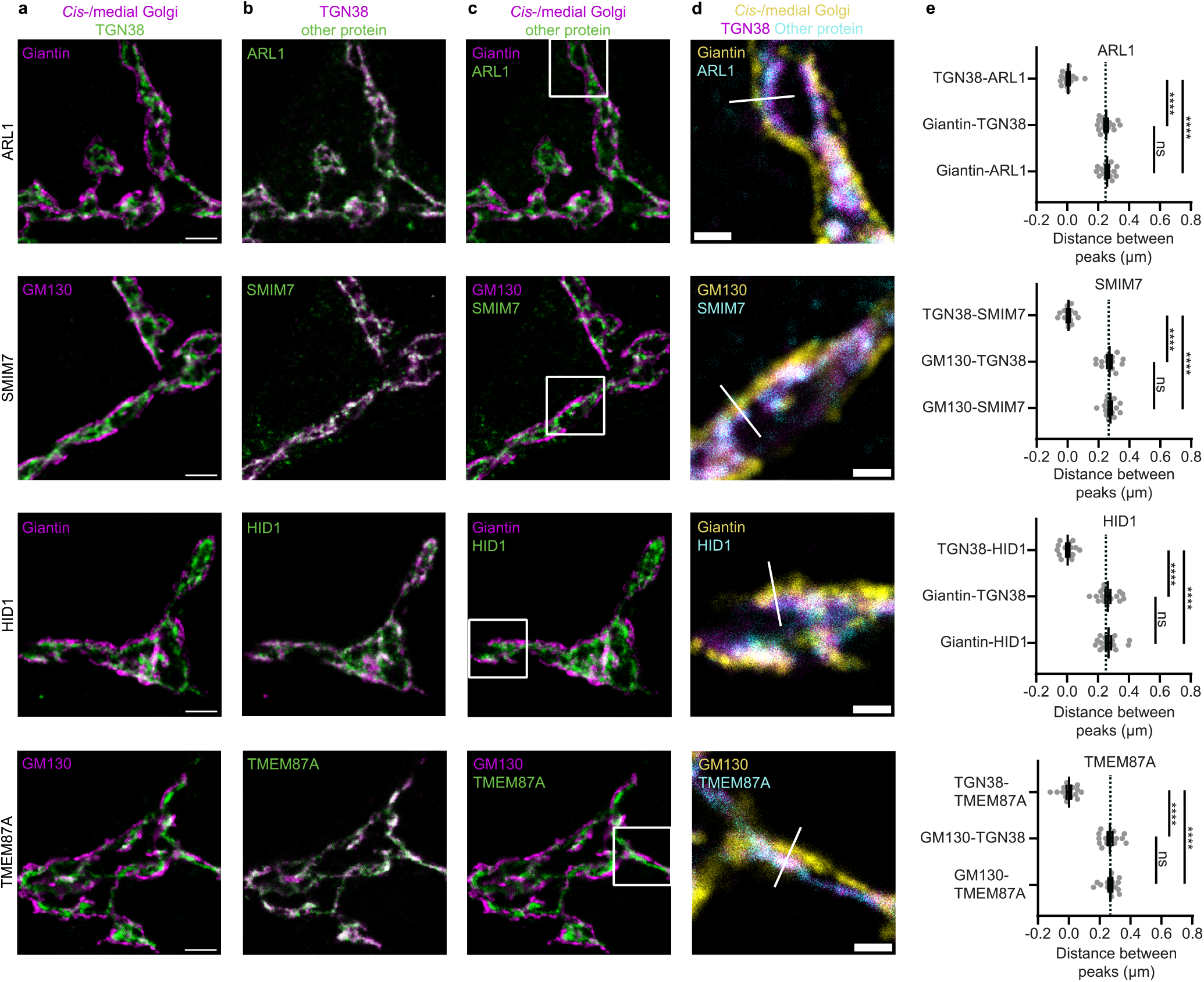
Several Golgi proteins overlap with TGN38. **a-c** Representative examples of neurons immunostained for a *cis*-/medial Golgi marker (GM130 or giantin), TGN38 and a protein of interest (ARL1, SMIM7, HID1 and TMEM87A). Images showing immunostaining of *cis*-/medial Golgi marker and TGN (**a**), TGN and protein of interest (**b**), *cis*-/medial Golgi and protein of interest (**c**). **d** Zoom from the white square area marked in **c**, showing all three stained proteins. White line is a representative example of a line drawn for intensity plotting. **e** Distance between intensity peaks of TGN38 and protein of interest, *cis*-/medial Golgi and TGN; and *cis*-/medial Golgi and protein of interest, measured by line scans. ARL1: (TGN38-ARL1: n = 15; Giantin-TGN38: n = 15; Giantin-ARL1: n = 15). One-way ANOVA with Tukey’s multiple comparisons test: TGN38-ARL1 vs Giantin-TGN38 and TGN38-ARL1 vs Giantin-ARL1: ****p < 0.0001, Giantin-TGN38 vs Giantin-ARL1: non-significant (ns). SMIM7: (TGN38-SMIM7: n = 15; GM130-TGN38: n = 15; GM130-SMIM7: n = 15). One-way ANOVA with Tukey’s multiple comparisons test: TGN38-SMIM7 vs GM130-TGN38 and TGN38-SMIM7 vs GM130-SMIM7: ****p < 0.0001, GM130-TGN38 vs GM130-SMIM7: non-significant (ns). HID1: (TGN38-HID1: n = 16; Giantin-TGN38: n = 16; Giantin-HID1: n = 16). One-way ANOVA with Tukey’s multiple comparisons test: TGN38-HID1 vs Giantin-TGN38 and TGN38-HID1 vs Giantin-HID1: ****p < 0.0001, Giantin-TGN38 vs Giantin-HID1: non-significant (ns). TMEM87A: (TGN38-TMEM87A: n = 15; GM130-TGN38: n = 15; GM130-TMEM87A: n = 15). One-way ANOVA with Tukey’s multiple comparisons test: TGN38-TMEM87A vs GM130-TGN38 and TGN38-TMEM87A vs GM130-TMEM87A: ****p < 0.0001, GM130-TGN38 vs GM130-TMEM87A: non-significant (ns). Dotted lines represent average distance between TGN38 and GM130 or TGN38 and giantin in the entire dataset. Bars show mean ± SEM. Detailed statistics are shown in Supplementary Table 1. Scale bar is 3 μm (**a**) and 1 μm (**d**).

**Fig. 4.**
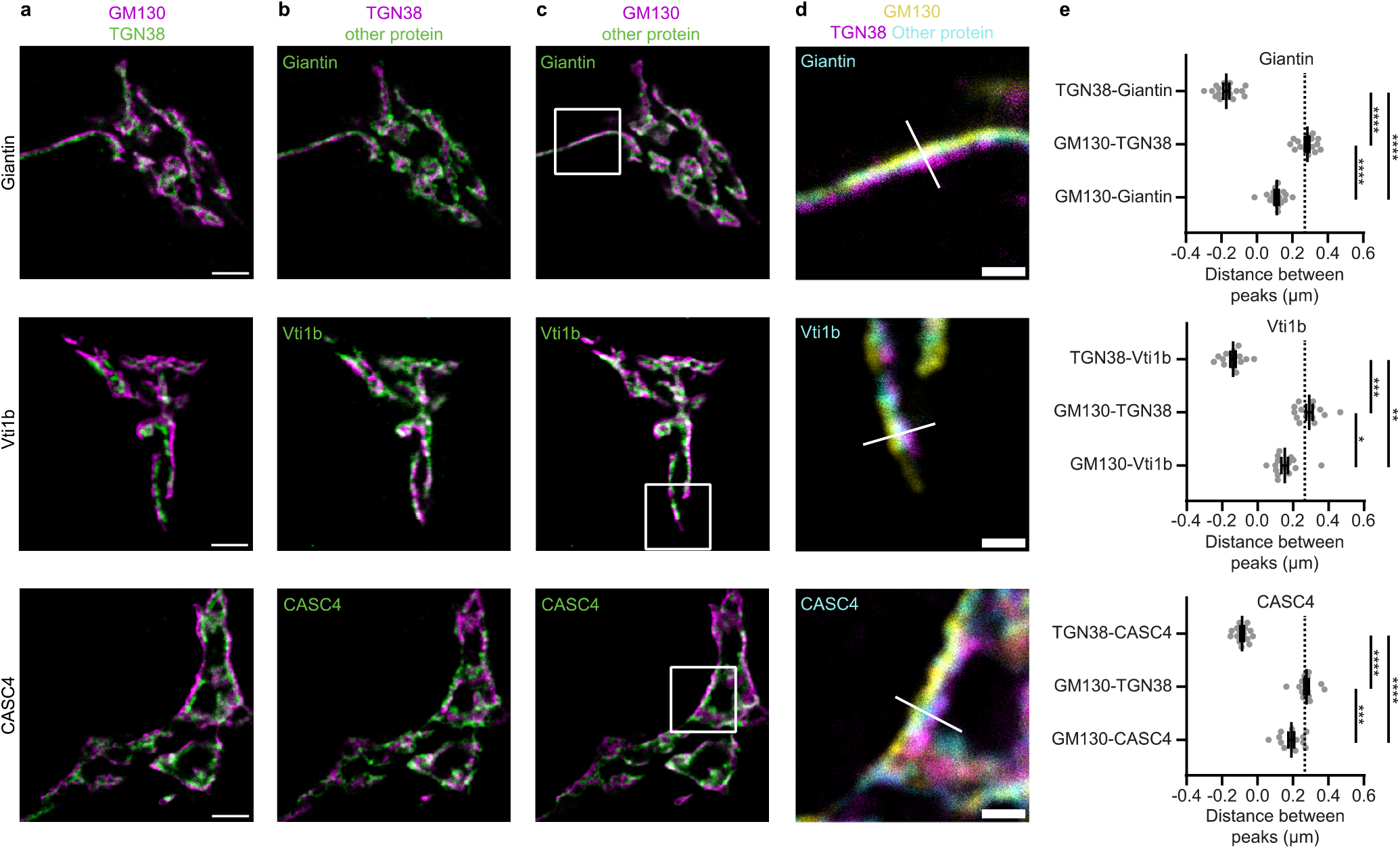
Several proteins localize upstream of TGN38. **a-c** Representative examples of neurons immunostained for *cis*-Golgi marker GM130, TGN38 and a protein of interest (Giantin, Vti1b and CASC4). Images showing immunostaining of *cis*-/medial Golgi marker and TGN (**a**), TGN and protein of interest (**b**), *cis*-/medial Golgi and protein of interest (**c**). **d** Zoom from the white square area marked in **c**, showing all three stained proteins. White line is a representative example of a line drawn for intensity plotting. **e** Distance between intensity peaks of TGN38 and protein of interest, *cis*-Golgi and TGN; and *cis*-Golgi and protein of interest, measured by line scans. Giantin: (TGN38-Giantin: n = 16; GM130-TGN38: n = 16; GM130-Giantin: n = 16). One-way ANOVA with Tukey’s multiple comparisons test: TGN38-Giantin vs GM130-TGN38, TGN38-Giantin vs GM130-Giantin and TGN38-GM130 vs GM130-Giantin: ****p < 0.0001. Vti1b: (TGN38-Vti1b: n = 15; GM130-TGN38: n = 15; GM130-Vti1b: n = 15). Kruskall-Wallis test with Dunn’s multiple comparisons test: TGN38-Vti1b vs GM130-TGN38: ****p < 0.0001, TGN38-Vti1b vs GM130-Vti1b: **p = 0.0025, TGN38-GM130 vs GM130-Vti1b: *p = 0.0201. CASC4: (TGN38-CASC4: n = 15; GM130-TGN38: n = 15; GM130-CASC4: n = 15). One-way ANOVA with Tukey’s multiple comparisons test: TGN38-CASC4 vs GM130-TGN38 and TGN38-CASC4 vs GM130-CASC4: ****p < 0.0001, TGN38-GM130 vs GM130-CASC4: ***p = 0.0001. Dotted lines represent average distance between TGN38 and GM130 in the entire dataset. Bars show mean ± SEM. Detailed statistics are shown in Supplementary Table 1. Scale bar is 3 μm (**a**) and 1 μm (**d**).

### Proteins overlapping with TGN38

ARL1, SMIM7, HID1 and TMEM87A overlapped with TGN38 (Fig. 3b). In addition, images in which *cis*-/medial Golgi and the protein of interest were shown were comparable to images in which *cis*-/medial Golgi and TGN were shown (Fig. 3a,c). In three-color images, only 2 layers were detected, because TGN38 and the protein of interest overlapped (Fig. 3d). The peak intensity coincided with the peak intensity of TGN38 and the distance between *cis*-/medial Golgi and proteins of interest was not different from the distance between *cis*-/medial Golgi and TGN38 (Fig. 3e). Supplementary figure 4 shows similar results for Golgin97, PRR36, TM9SF3 and Vti1a. However, a large variation in localization of Vti1a was observed and peak intensity of Vti1a staining did not overlap as perfectly with TGN38 as the other seven TGN proteins. Taken together, ARL1, SMIM7, HID1, TMEM87A, Golgin97, PRR36, TM9SF3 and Vti1a localized close to TGN38.

### Proteins upstream from TGN38

Giantin, Vti1b and CASC4 did not colocalize with TGN38 or GM130 (Fig. 4b,c), but the peak intensity of their staining localized in between GM130 and TGN38 (Fig. 4d). This was also shown by a positive distance from GM130 and a negative distance from TGN38 (Fig. 4e). Thus, giantin, Vti1b and CASC4 were upstream from TGN38. Figure 5 provides an overview of all proteins stained in this project and their distance from TGN38.

**Fig. 5.**
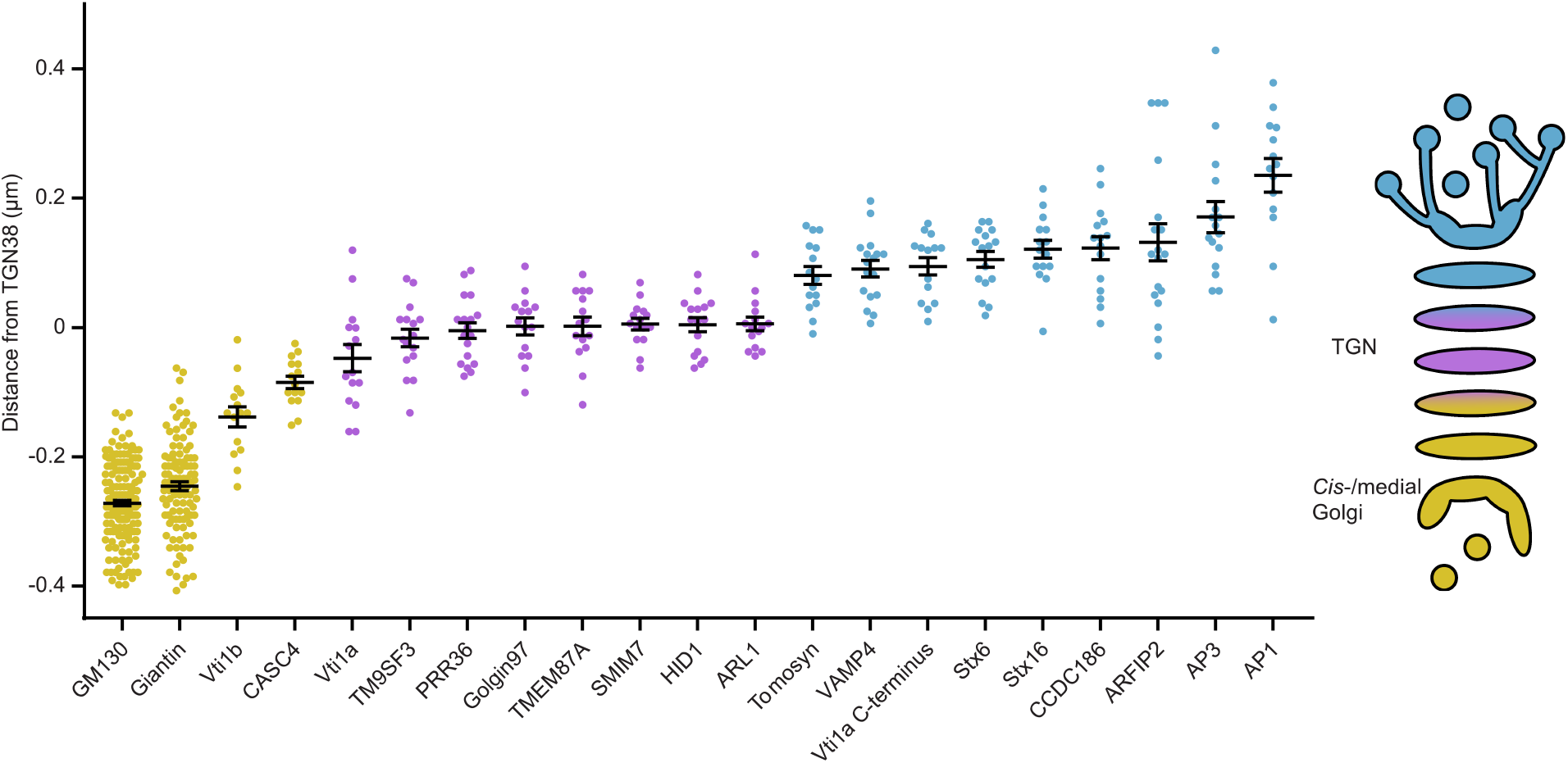
Overview of Golgi protein distribution relative to TGN38. Proteins upstream of TGN38 are shown in yellow, proteins overlapping with TGN38 in magenta and proteins downstream of TGN38 in cyan. Bars show mean ± SEM.

## Discussion

In this study, we analyzed the distribution of 21 proteins in the Golgi apparatus of primary mouse neurons. These proteins were categorized into three groups based on their relative localization along the Golgi axis: proteins at upstream of TGN38 (*cis*-/medial Golgi), proteins overlapping with TGN38 (TGN) and proteins downstream of TGN38. The proteins downstream from TGN38 were localized at varying distances from TGN38, suggesting they are in different sub-compartments of the TGN. Seven proteins overlapped perfectly with TGN38 and four proteins were upstream of TGN38, localizing to the *cis*-/medial Golgi. The relative localization of proteins in the Golgi is consistent with previous models made by Tie et al. [4,5,9]. The most striking difference is the number of proteins downstream of TGN38 in our model.

Proteins downstream of TGN38 likely localize to sub-compartments of the TGN, consistent with their function. AP1 and AP3, for example, are adapter proteins involved in vesicle budding [10,11], which happens on the outer borders of the TGN. The distance between AP1 and TGN38 is almost as large as between TGN38 and giantin. Similarly, ARFIP2 senses membrane curvature and induces membrane tubulation [12,13]. We did observe large variation in ARFIP2 localization, perhaps indicating different steps and/or locations of membrane tubulation. CCDC186 plays a role in dense core vesicle biogenesis and was previously described to be near to, but not overlapping with TGN38 [14]. This conclusion is consistent with our data.

In decreasing distance from TGN38 we identified Stx16, Stx6, Vti1a and VAMP4. These proteins form a SNARE complex involved in retrograde trafficking from early endosomes to the TGN [15,16]. It was previously shown that the formation and docking of vesicles containing mannose 6-phosphate receptors (MPRs) happens at distinct domains of the TGN [17]. Similarly, the TGN of fission yeast was divided into early and late TGN: early TGN for cargo reception and late TGN for vesicle formation [18]. In agreement with this, we observed that proteins involved in vesicle budding, such as AP1 and AP3, are at a different part of the TGN than proteins involved in fusion of retrograde vesicles, such as Stx6 and Stx16. The two antibodies against Vti1a used in the current study, showed a different localization. Previously, immunoblotting using an antibody against the C-terminus of Vti1a resulted in one band of 14kDa, while an antibody raised against a bigger part of Vti1a yielded multiple bands [19]. This could explain why the C-terminal antibody results in a more precise localization in our experiments, while localization of the other antibody shows a broader distribution. Tomosyn is mainly known for inhibiting priming of synaptic and dense core vesicles by forming nonfusogenic SNARE complexes [20–22] and localization to the TGN has only been described in Arabidopsis protoplasts [23]. We observed high intensity of tomosyn staining downstream of TGN38, suggesting that tomosyn also has a function at the TGN.

Another group of proteins overlapped with TGN38. HID1 is thought to play a role in protein sorting by regulating TGN pH [24] and was previously shown to colocalize with TGN38 [25]. Arl1 recruits golgin97, a tether protein involved in retrograde trafficking from endosomes to TGN [26]. The vesicles tethered by golgin97 contain both TGN38 and TMEM87A [27]. This is in agreement with our data, showing that ARL1, golgin97 and TMEM87A overlap with TGN38. The exact function of SMIM7, PRR36, TM9SF3 and CASC4 is unknown. Consistent with its function in the fusion of late endosomes [28], Vti1b was described to associate with endosomes [29]. In addition, it was found in the TGN [29,30]. Vti1p, the yeast homolog of Vti1a and Vti1b, is also involved in intra-Golgi trafficking [31]. Since loss of Vti1a and Vti1b dramatically affects Golgi organization while single knockout of either one of these proteins does not affect the Golgi in mammalian neurons [32,33], it is plausible that Vti1a and Vti1b also play a role in intra-Golgi trafficking in mouse neurons. GM130, Giantin and TGN38 are commonly used markers for *cis*-, *cis*-/medial Golgi and TGN, respectively [34–36]. Taken together, the relative localization of the analyzed proteins is consistent with previously described localization and their proposed functions.

## Methods

### Laboratory animals and primary cultures

Animal experiments were approved by the animal ethical committee of the VU University/VU University Medical Centre (license number: FGA 11-03). Animals were bred and housed according to Institutional and Dutch governmental guidelines. All experiments were conducted in compliance with ARRIVE guidelines. Embryonic day (E) 18.5 C57BL/6 mouse embryos were used for primary hippocampal culture. For this study, three wildtype pups were used. Mouse hippocampi were dissected in Hanks’ balanced salt solution (Sigma), supplemented with 10mM HEPES (Gibco) and were digested with 0.25% trypsin (Gibco) in Hanks’ + HEPES for 15 min at 37°C. Hippocampi were washed three times with Hanks’ + HEPES, once with DMEM complete (DMEM + Glutamax (Gibco), supplemented with 10% FCS (Gibco), 1% NEAA (Sigma) and 1% penicillin/streptomycin (Sigma)) and triturated with fire-polished Pasteur pipettes. Dissociated cells were spun down and resuspended in Neurobasal medium (Gibco) supplemented with 2% B-27 (Gibco), 1.8% HEPES, 0.25% Glutamax (Gibco) and 0.1% penicillin/streptomycin. Continental cultures were created by plating neurons at 25K/well. Neurons were seeded on pre-grown rat glia on 18mm glass coverslips in 12 well plates.

### Immunocytochemistry

Neurons were fixed at DIV14 in freshly prepared 3.7% paraformaldehyde (EMS) for 10 min at room temperature and permeabilized with 0.1% Triton X-100 (Fisher Scientific) for 10 min. Incubation with primary and secondary antibodies was done at room temperature for 1 h. To combine donkey anti-sheep and goat antibodies, secondary antibody incubation was split into two steps to prevent cross-reaction: donkey anti-sheep antibodies were incubated for 1h and after thorough washing and blocking with 5% normal goat serum (Life Technologies) for 30 min, goat secondary antibodies were incubated for 1h. All solutions were in PBS (composition in mM: 137 NaCl, 2.7 KCl, 10 Na2HPO4, 1.8 KH2PO4; pH= 7.4). Coverslips were mounted in Mowiol (Sigma).

#### Primary antibodies used for immunocytochemistry

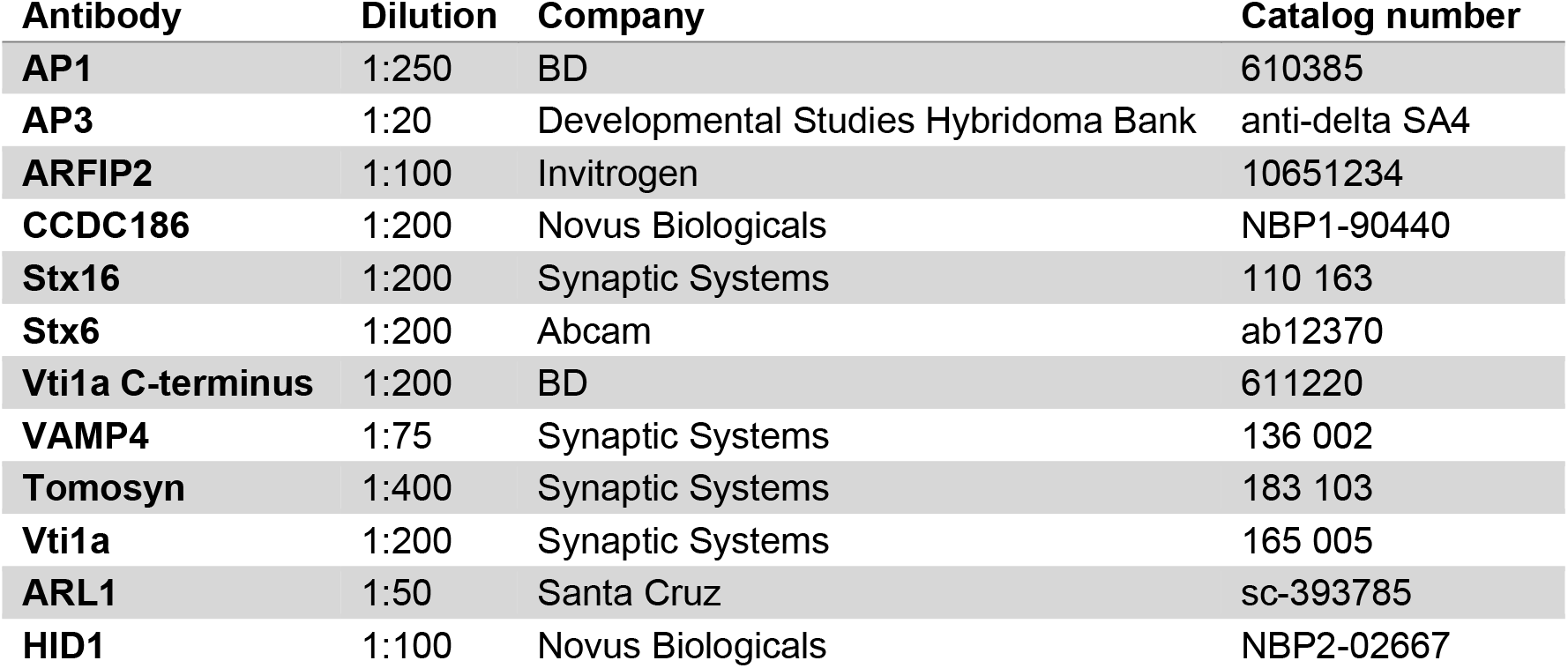
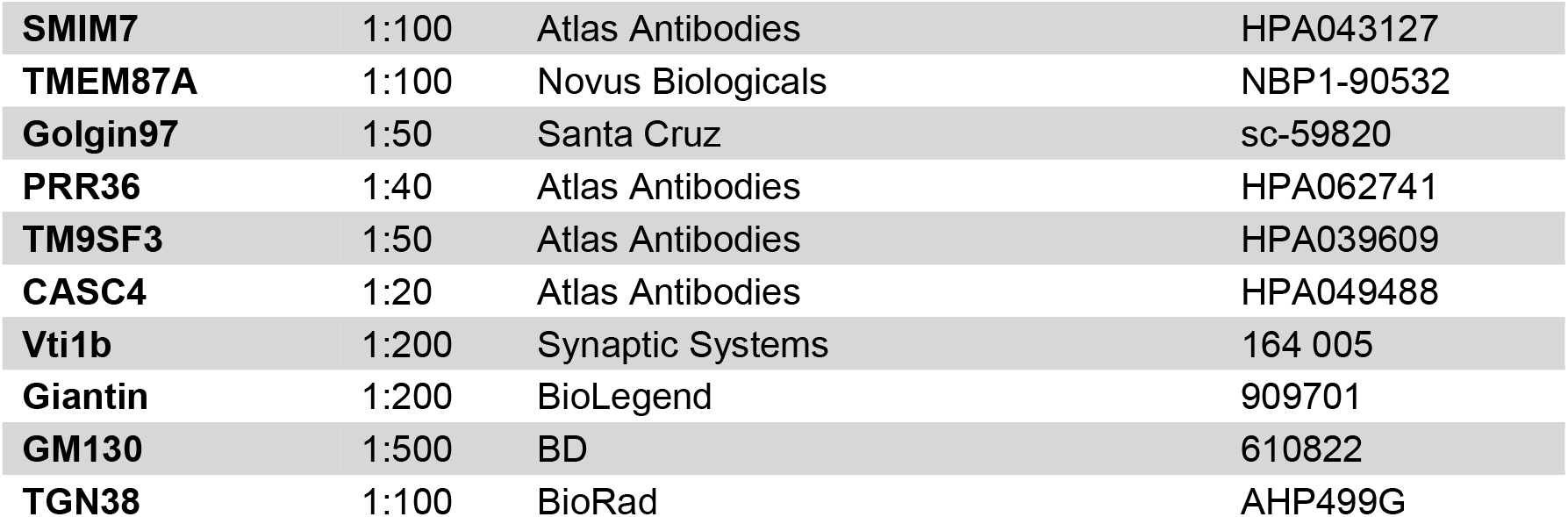

Secondary antibodies were AlexaFluor-conjugated (1:100, Invitrogen) or STAR580- or STAR635p-conjugated (1:100, Abberior).

Confocal imaging was performed on a Leica TCS SP8 STED 3x microscope (Leica Microsystems) with LAS X acquisition software, using a 100x oil objective (NA=1.4). A pulsed white light laser was used to excite the samples. The signal was detected using a gated hybrid detector (HyD) (Leica Microsystems) in photon-counting mode.

Images were analyzed in Fiji software [37]. The BAR plugin was used to plot multichannel profiles in order to analyze the distance between peaks of proteins in the Golgi [38]. We chose to plot distance relative to TGN38, because the TGN38 sheep antibody we used could be combined with any antibody species for the protein of interest. This is in contrast to for example rabbit giantin which cannot be combined with rabbit CCDC186.

### Statistical analysis

Statistical analysis and graphing were performed using GraphPad Prism 7. Shapiro-Wilk was used to test distribution normality. When assumption of normality was met, parametric tests were used: two-tailed *t*-test or ANOVA (with Tukey’s multiple comparisons test). Otherwise, non-parametric tests were used: two-tailed Mann Whitney or Kruskall-Wallis (with Dunn’s multiple comparisons test). Significance was accepted at p < 0.05. Each data point represents one cell and bars show mean ± SEM. All data and statistical tests used are summarized in Supplementary Table 1. No sample size calculation was performed. No randomization or blinding was performed. Neurons were excluded from analysis if neuronal Golgi could not be distinguished, for example because it was overlapping with glial Golgi.

## Data availability

The data generated and analyzed during the current study are included in this published article and its supplementary information files. Datasets are available from the corresponding author on request.

## Acknowledgements

We thank Lisa Laan, Ingrid Saarloos and Desiree Schut for producing glia and for primary culture assistance and Joke Wortel for animal breeding. We acknowledge the Microscopy and Cytometry Core Facility at the Amsterdam UMC – Location VUmc for providing assistance in microscopy. This work is supported by a European Research Council Advanced Grant (322966) of the European Union (to M.V.).

## Author contributions

D.V.B. performed experiments and analyzed the data. D.V.B., R.F.T. and M.V. designed the research and wrote the paper.

