## Supplementary information for "Mapping localization of 21 endogenous proteins in the Golgi apparatus of rodent neurons"

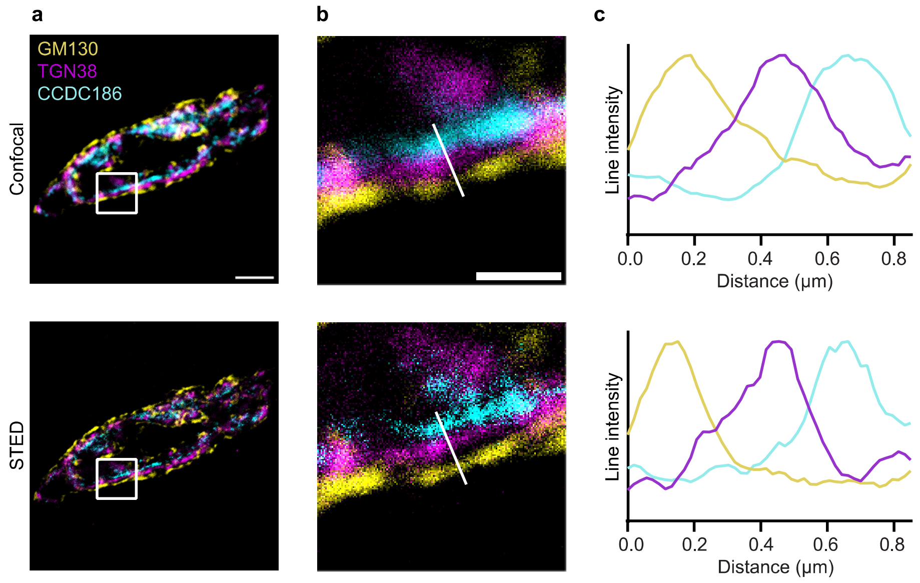


**Supplementary Fig. 1** Comparison of confocal and STED microscopy. **a** Confocal and STED image of a neuron stained for GM130 (*cis*-Golgi), TGN38 (TGN) and CCDC186. **b** Zoom from the white square area marked in **a**. **c** Line intensity profile along white line in **b**. Intensities were normalized and smoothed before plotting. Scale bar is 3 µm (**a**) and 1 µm (**b**).

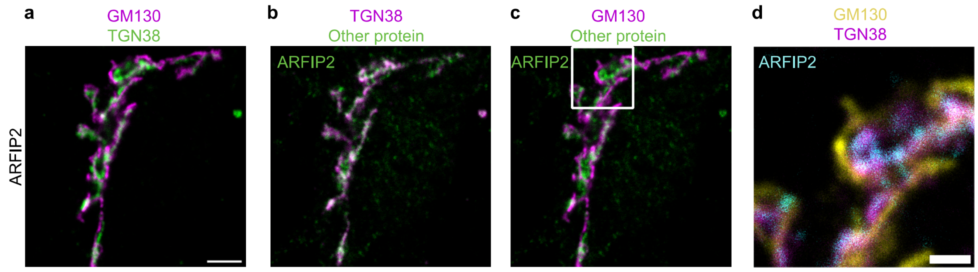


**Supplementary Fig. 2** Alternative localization of ARFIP2. **a-c** Representative examples of neurons immunostained for *cis*-Golgi marker GM130, TGN38 and ARFIP2. Images showing immunostaining of *cis*-Golgi and TGN (**a**), TGN and ARFIP2 (**b**), *cis*-Golgi and ARFIP2 (**c**). **d** Zoom from the white square area marked in **c**, showing all three stained proteins. Scale bar is 3 µm (**a**) and 1 µm (**d**).

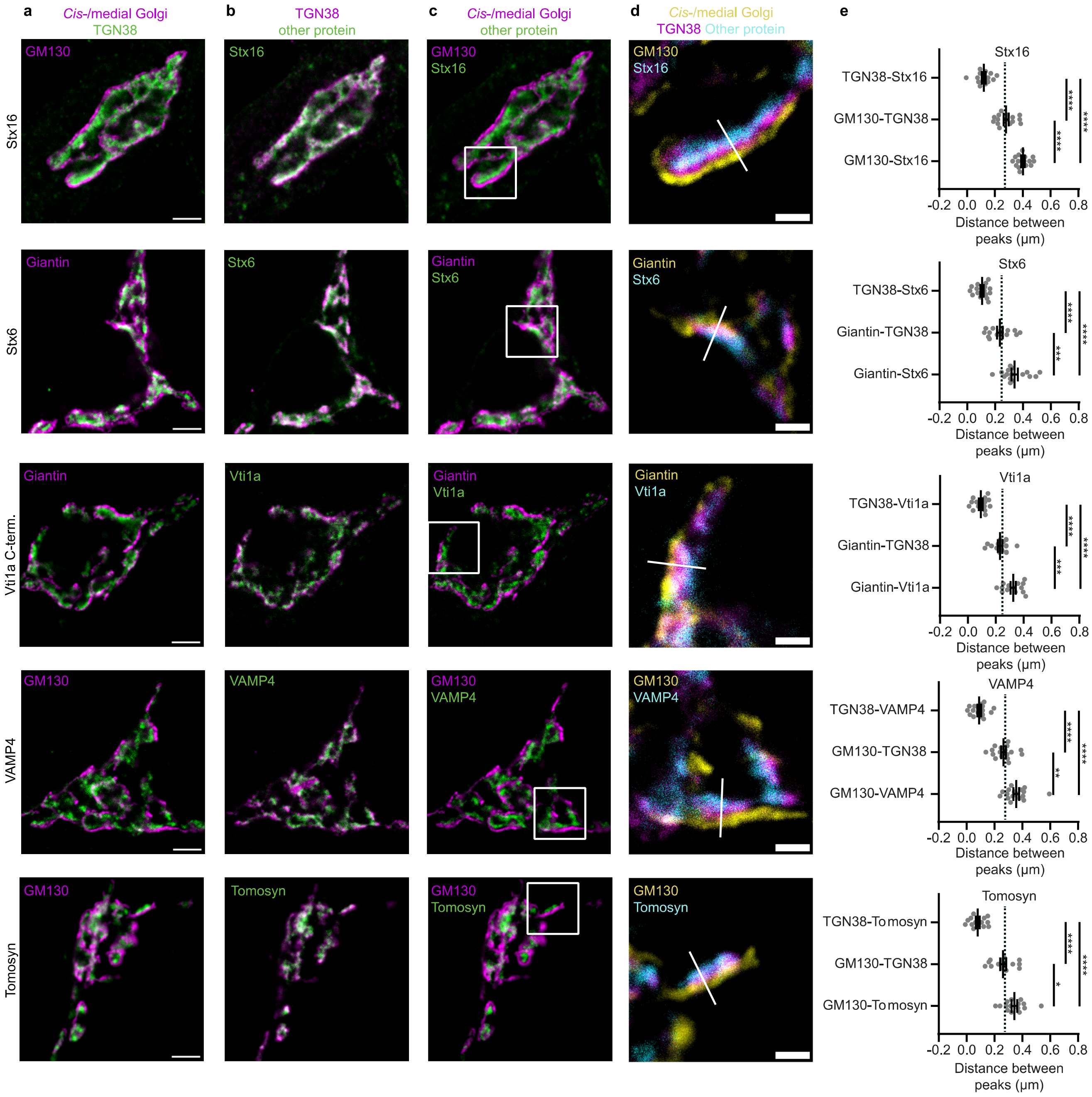


**Supplementary Fig. 3** Several Golgi proteins localize downstream of TGN38. **a-c** Representative examples of neurons immunostained for a *cis*-/medial Golgi marker (GM130 or giantin), TGN38 and a protein of interest (Stx16, Stx6, Vti1a C-terminus, VAMP4 and Tomosyn). Images showing immunostaining of *cis*-/medial Golgi marker and TGN (**a**), TGN and protein of interest (**b**), *cis*-/medial Golgi and protein of interest (**c**). **d** Zoom from the white square area marked in **c**, showing all three stained proteins. White line is a representative example of a line drawn for intensity plotting. **e** Distance between intensity peaks of TGN38 and protein of interest, *cis*-/medial Golgi and TGN; and *cis*-/medial Golgi and protein of interest, measured by line scans. Stx16: (TGN38-Stx16: n = 15; GM130-TGN38: n = 15; GM130-Stx16: n = 15). One-way ANOVA with Tukey's multiple comparisons test: TGN38-Stx16 vs GM130-TGN38, TGN38-Stx16 vs GM130-Stx16 and GM130-TGN38 vs GM130-Stx16: ****p < 0.0001. Stx6: (TGN38-Stx6: n = 16; Giantin-TGN38: n = 16; Giantin-Stx6: n = 16). One-way ANOVA with Tukey's multiple comparisons test: TGN38-Stx6 vs Giantin-TGN38 and TGN38-Stx6 vs Giantin-Stx6: ****p < 0.0001, Giantin-TGN38 vs Giantin-Stx6: ***p = 0.0008. Vti1a C-term.: (TGN38-Vti1a C-term.: n = 14; Giantin-TGN38: n = 14; Giantin-Vti1a C-term.: n = 14). One-way ANOVA with Tukey's multiple comparisons test: TGN38-Vti1a C-term. vs Giantin-TGN38 and TGN38-Vti1a C-term. vs Giantin-Vti1a C-term.: ****p < 0.0001, Giantin-TGN38 vs Giantin-Vti1a C-term.: ***p = 0.0005. VAMP4: (TGN38-VAMP4: n = 17; GM130-TGN38: n = 17; GM130-VAMP4: n = 17). One-way ANOVA with Tukey's multiple comparisons test: TGN38-VAMP4 vs GM130-TGN38 and TGN38-VAMP4 vs GM130-VAMP4: ****p < 0.0001, and GM130-TGN38 vs GM130-VAMP4: **p = 0.0011. Tomosyn: (TGN38-Tomosyn: n = 15; GM130-TGN38: n = 15; GM130-Tomosyn: n = 15). One-way ANOVA with Tukey's multiple comparisons test: TGN38-Tomosyn vs GM130-TGN38 and TGN38-Tomosyn vs GM130-Tomosyn: ****p < 0.0001, GM130-TGN38 vs GM130-Tomosyn: *p = 0.0109. Dotted lines represent average distance between TGN38 and GM130 or TGN38 and giantin in the entire dataset. Bars show mean ± SEM. Detailed statistics are shown in Supplementary Table 1. Scale bar is 3 µm (**a**) and 1 µm (**d**).

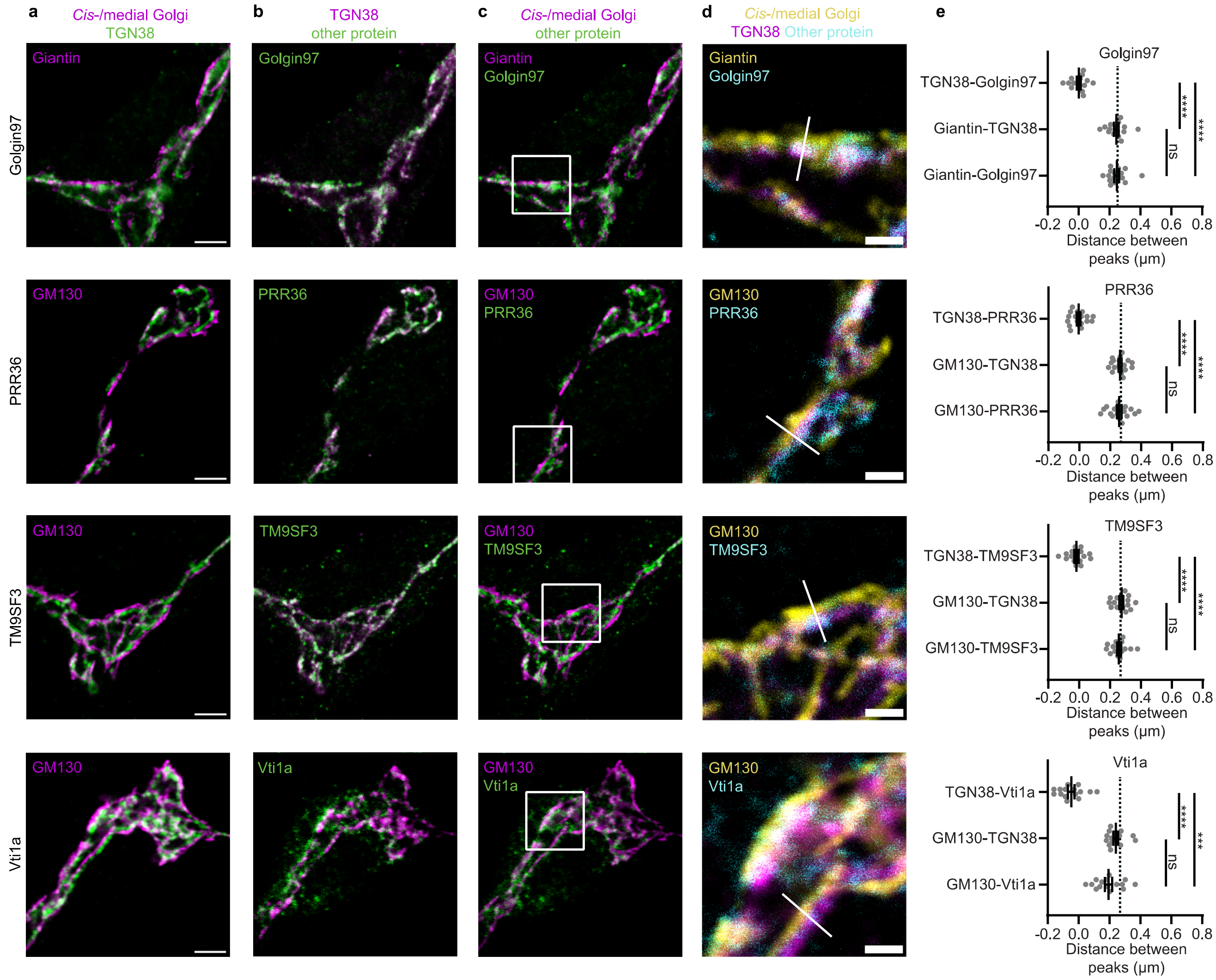


**Supplementary Fig. 4** Several Golgi proteins overlap with TGN38. **a-c** Representative examples of neurons immunostained for a *cis*-/medial Golgi marker (GM130 or giantin), TGN38 and a protein of interest (TMEM87A, Golgin97, PRR36 and TM9SF3). Images showing immunostaining of *cis*-/medial Golgi marker and TGN (**a**), TGN and protein of interest (**b**), *cis*-/medial Golgi and protein of interest (**c**). **d** Zoom from the white square area marked in **c**, showing all three stained proteins. White line is a representative example of a line drawn for intensity plotting. **e** Distance between intensity peaks of TGN38 and protein of interest, *cis*-/medial Golgi and TGN; and *cis*-/medial Golgi and protein of interest, measured by line scans. Golgin97: (TGN38-Golgin97: n = 15; Giantin-TGN38: n = 15; Giantin-Golgin97: n = 15). One-way ANOVA with Tukey's multiple comparisons test: TGN38-Golgin97 vs Giantin-TGN38 and TGN38-Golgin97 vs Giantin-Golgin97: ****$p < 0.0001$, Giantin-TGN38 vs Giantin-Golgin97: non-significant (ns). PRR36: (TGN38-PRR36: n = 18; GM130-TGN38: n = 18; GM130-PRR36: n = 18). One-way ANOVA with Tukey's multiple comparisons test: TGN38-PRR36 vs GM130-TGN38 and TGN38-PRR36 vs GM130-PRR36: ****$p < 0.0001$, GM130-TGN38 vs GM130-PRR36: non-significant (ns). TM9SF3: (TGN38-TM9SF3: n = 16; GM130-TGN38: n = 16; GM130-TM9SF3: n = 16). One-way ANOVA with Tukey's multiple comparisons test: TGN38-TM9SF3 vs GM130-TGN38 and TGN38-TM9SF3 vs GM130-TM9SF3: ****$p < 0.0001$, GM130-TGN38 vs GM130-TM9SF3: non-significant (ns). Vti1a: (TGN38-Vti1a: n = 15; GM130-TGN38: n = 15; GM130-Vti1a: n = 15). Kruskall-Wallis test with Dunn's multiple comparisons test: TGN38-Vti1a vs GM130-TGN38: ****$p < 0.0001$, TGN38-Vti1a vs GM130-Vti1a: ***$p = 0.0002$, GM130-TGN38 vs GM130-Vti1a: non-significant (ns). Dotted lines represent average distance between TGN38 and GM130 or TGN38 and giantin in the entire dataset. Bars show mean ± SEM. Detailed statistics are shown in Supplementary Table 1. Scale bar is 3 µm (**a**) and 1 µm (**d**).

**Supplementary Table 1: Data and statistics.** Overview of all data and statistics for each figure. Dataset, condition, average and SEM, the number of independent cells (n), the p-values and statistical tests used are indicated. Statistical tests were two-tailed and used α = 0.05. *p < 0.05; **p < 0.01; ***p < 0.001; ****p < 0.0001. For one-way ANOVA the p-values are also indicated.

| Dataset | Condition | Value (Mean ± SEM) | n | p-value | Statistical test |
|---|---|---|---|---|---|
| AP1 Figure 2e | (1) TGN38-AP1 | 0.236 ± 0.0262 | 14 | ns, p = 0.6031: 1 versus 2, ****p <0.0001: 1 versus 3, **p = 0.0036: 2 versus 3 | Kruskal-Wallis test (****p < 0.0001) with Dunn's multiple comparisons test |
| | (2) Giantin-TGN38 | 0.295 ± 0.0176 | 14 | | |
| | (3) Giantin-AP1 | 0.531 ± 0.0338 | 14 | | |
| AP3 Figure 2e | (1) TGN38-AP3 | 0.171 ± 0.0243 | 16 | ns, p = 0.0940: 1 versus 2, ****p <0.0001: 1 versus 3, **p = 0.0042: 2 versus 3 | Kruskal-Wallis test (****p < 0.0001) with Dunn's multiple comparisons test |
| | (2) Giantin-TGN38 | 0.262 ± 0.0149 | 16 | | |
| | (3) Giantin-AP3 | 0.433 ± 0.0293 | 16 | | |
| ARFIP2 Figure 2e | (1) TGN38-ARFIP2 | 0.132 ± 0.0289 | 18 | **p = 0.0018: 1 versus 2, ****p <0.0001: 1 versus 3, ***p = 0.0009: 2 versus 3 | One-way ANOVA (****p < 0.0001) with Tukey's multiple comparisons test |
| | (2) GM130-TGN38 | 0.256 ± 0.0125 | 18 | | |
| | (3) GM130-ARFIP2 | 0.388 ± 0.0272 | 18 | | |
| CCDC186 Figure 2e | (1) TGN38-CCDC186 | 0.123 ± 0.0178 | 15 | ****p < 0.0001: 1 versus 2, 1 versus 3, ***p = 0.0004: 2 versus 3 | One-way ANOVA (****p < 0.0001) with Tukey's multiple comparisons test |
| | (2) GM130-TGN38 | 0.29 ± 0.0163 | 15 | | |
| | (3) GM130-CCDC186 | 0.412 ± 0.0262 | 15 | | |
| ARL1 Figure 3e | (1) TGN38-ARL1 | 0.00589 ± 0.0107 | 15 | ****p < 0.0001: 1 versus 2, 1 versus 3, ns, p = 0.9277: 2 versus 3 | One-way ANOVA (****p < 0.0001) with Tukey's multiple comparisons test |
| | (2) Giantin-TGN38 | 0.252 ± 0.0122 | 15 | | |
| | (3) Giantin-ARL1 | 0.258 ± 0.109 | 15 | | |
| SMIM7 Figure 3e | (1) TGN38-SMIM7 | 0.00526 ± 0.00888 | 15 | ****p < 0.0001: 1 versus 2, 1 versus 3, ns, p = 0.9422: 2 versus 3 | One-way ANOVA (****p < 0.0001) with Tukey's multiple comparisons test |
| | (2) GM130-TGN38 | 0.271 ± 0.014 | 15 | | |
| | (3) GM130-SMIM7 | 0.277 ± 0.104 | 15 | | |
| HID1 Figure 3e | (1) TGN38-HID1 | 0.00454 ± 0.011 | 16 | ****p < 0.0001: 1 versus 2, 1 versus 3, ns, p = 0.9744: 2 versus 3 | One-way ANOVA (****p < 0.0001) with Tukey's multiple comparisons test |
| | (2) Giantin-TGN38 | 0.264 ± 0.0164 | 16 | | |
| | (3) Giantin-HID1 | 0.269 ± 0.0163 | 16 | | |
| TMEM87A Figure 3e | (1) TGN38-TMEM87A | 0.0021 ± 0.0142 | 15 | ****p < 0.0001: 1 versus 2, 1 versus 3, ns, p = 0.9942: 2 versus 3 | One-way ANOVA (****p < 0.0001) with Tukey's multiple comparisons test |
| | (2) GM130-TGN38 | 0.267 ± 0.0157 | 15 | | |
| | (3) GM130-TMEM87A | 0.27 ± 0.0134 | 15 | | |
| Giantin Figure 4e | (1) TGN38-Giantin | -0.172 ± 0.0171 | 16 | ****p < 0.0001: 1 versus 2, 1 versus 3, 2 versus 3 | One-way ANOVA (****p < 0.0001) with Tukey's multiple comparisons test |
| | (2) GM130-TGN38 | 0.285 ± 0.0131 | 16 | | |
| | (3) GM130-Giantin | 0.113 ± 0.0123 | 16 | | |
| Vti1b Figure 4e | (1) TGN38-Vti1b | -0.138 ± 0.0155 | 15 | ****p < 0.0001: 1 versus 2, **p = 0.0025: 1 versus 3, *p = 0.0201: 2 versus 3 | Kruskal-Wallis test (****p < 0.0001) with Dunn's multiple comparisons test |
| | (2) GM130-TGN38 | 0.292 ± 0.0185 | 15 | | |
| | (3) GM130-Vti1b | 0.154 ± 0.0188 | 15 | | |
| CASC4 Figure 4e | (1) TGN38-CASC4 | -0.0854 ± 0.00972 | 15 | ****p < 0.0001: 1 versus 2, 1 versus 3, ns, p = 0.9610: 2 versus 3 | One-way ANOVA (****p < 0.0001) with Tukey's multiple comparisons test |
| | (2) GM130-TGN38 | 0.278 ± 0.0127 | 15 | | |
| | (3) GM130-CASC4 | 0.192 ± 0.016 | 15 | | |
| Distance from TGN38 Figure 5 | GM130 | -0.272 ± 0.00442 | 175 | | |
| | Giantin | -0.246 ± 0.00693 | 108 | | |
| | Vti1b | -0.138 ± 0.0155 | 15 | | |
| | CASC4 | -0.0854 ± 0.00972 | 15 | | |
| | Vti1a | -0.0473 ± 0.0209 | 15 | | |

| | | | | | |
|---|---|---|---|---|---|
| | TM9SF3 | -0.0162 ± 0.0138 | 16 | | |
| | PRR36 | -0.00491 ± 0.012 | 18 | | |
| | Golgin97 | 0.00189 ± 0.0131 | 15 | | |
| | TMEM87A | 0.0021 ± 0.0142 | 15 | | |
| | SMIM7 | 0.00526 ± 0.00888 | 15 | | |
| | HID1 | 0.00454 ± 0.011 | 16 | | |
| | ARL1 | 0.00589 ± 0.0107 | 15 | | |
| | Tomosyn | 0.0808 ± 0.0138 | 15 | | |
| | VAMP4 | 0.0908 ± 0.0127 | 17 | | |
| | Vti1a C-terminus | 0.0947 ± 0.0136 | 14 | | |
| | Stx6 | 0.106 ± 0.0121 | 16 | | |
| | Stx16 | 0.121 ± 0.0138 | 15 | | |
| | CCDC186 | 0.123 ± 0.0178 | 15 | | |
| | ARFIP2 | 0.132 ± 0.0289 | 18 | | |
| | AP3 | 0.171 ± 0.0243 | 16 | | |
| | AP1 | 0.236 ± 0.0262 | 14 | | |
| Stx16 Sup. Figure 2e | (1) TGN38-Stx16 | 0.121 ± 0.0138 | 15 | ****p < 0.0001: 1 versus 2, 1 versus 3, 2 versus 3 | One-way ANOVA (****p < 0.0001) with Tukey's multiple comparisons test |
| | (2) GM130-TGN38 | 0.281 ± 0.0175 | 15 | | |
| | (3) GM130-Stx16 | 0.402 ± 0.0123 | 15 | | |
| Stx6 Sup. Figure 2e | (1) TGN38-Stx6 | 0.106 ± 0.0121 | 16 | ****p < 0.0001: 1 versus 2, 1 versus 3, ***p = 0.0008: 2 versus 3 | One-way ANOVA (****p < 0.0001) with Tukey's multiple comparisons test |
| | (2) Giantin-TGN38 | 0.235 ± 0.02 | 16 | | |
| | (3) Giantin-Stx6 | 0.34 ± 0.0231 | 16 | | |
| Vti1a C-terminus Sup. Figure 3e | (1) TGN38-Vti1a | 0.0947 ± 0.0136 | 14 | ****p < 0.0001: 1 versus 2, 1 versus 3, ***p = 0.0005: 2 versus 3 | One-way ANOVA (****p < 0.0001) with Tukey's multiple comparisons test |
| | (2) Giantin-TGN38 | 0.232 ± 0.0159 | 14 | | |
| | (3) Giantin-Vti1a | 0.326 ± 0.0188 | 14 | | |
| VAMP4 Sup. Figure 3e | (1) TGN38-VAMP4 | 0.0908 ± 0.0127 | 17 | ****p < 0.0001: 1 versus 2, 1 versus 3, **p = 0.0011: 2 versus 3 | One-way ANOVA (****p < 0.0001) with Tukey's multiple comparisons test |
| | (2) GM130-TGN38 | 0.266 ± 0.017 | 17 | | |
| | (3) GM130-VAMP4 | 0.357 ± 0.0198 | 17 | | |
| Tomosyn Sup. Figure 3e | (1) TGN38-Tomosyn | 0.0808 ± 0.0138 | 15 | ****p < 0.0001: 1 versus 2, 1 versus 3, *p = 0.0109: 2 versus 3 | One-way ANOVA (****p < 0.0001) with Tukey's multiple comparisons test |
| | (2) GM130-TGN38 | 0.262 ± 0.0207 | 15 | | |
| | (3) GM130-Tomosyn | 0.343 ± 0.0209 | 15 | | |
| Golgin97 Sup. Figure 4e | (1) TGN38-Golgin97 | 0.00189 ± 0.0131 | 15 | ****p < 0.0001: 1 versus 2, 1 versus 3, ns, p = 0.9955: 2 versus 3 | One-way ANOVA (****p < 0.0001) with Tukey's multiple comparisons test |
| | (2) Giantin-TGN38 | 0.246 ± 0.0154 | 15 | | |
| | (3) Giantin-Golgin97 | 0.247 ± 0.0159 | 15 | | |
| PRR36 Sup. Figure 4e | (1) TGN38-PRR36 | -0.00491 ± 0.012 | 18 | ****p < 0.0001: 1 versus 2, 1 versus 3, ns, p = 0.9610: 2 versus 3 | One-way ANOVA (****p < 0.0001) with Tukey's multiple comparisons test |
| | (2) GM130-TGN38 | 0.263 ± 0.0105 | 18 | | |
| | (3) GM130-PRR36 | 0.258 ± 0.0157 | 18 | | |
| TM9SF3 Sup. Figure 4e | (1) TGN38-TM9SF3 | -0.0162 ± 0.0138 | 16 | ****p < 0.0001: 1 versus 2, 1 versus 3, ns, p = 0.6640: 2 versus 3 | One-way ANOVA (****p < 0.0001) with Tukey's multiple comparisons test |
| | (2) GM130-TGN38 | 0.275 ± 0.0127 | 16 | | |
| | (3) GM130-TM9SF3 | 0.259 ± 0.0131 | 16 | | |
| Vti1a Sup. Figure 4e | (1) TGN38-Vti1a | -0.0473 ± 0.0209 | 15 | ****p < 0.0001: 1 versus 2, ***p = 0.0002: 1 versus 3, ns, p = 0.6479: 2 versus 3 | Kruskal-Wallis test (****p < 0.0001) with Dunn's multiple comparisons test |
| | (2) GM130-TGN38 | 0.241 ± 0.0148 | 15 | | |
| | (3) GM130-Vti1a | 0.194 ± 0.0232 | 15 | | |
